# High-Resolution Melting Analysis of Chloroplast Markers for Species Authentication and Fraud Detection in Commercial Açaí and Juçara Products

**DOI:** 10.64898/2026.05.01.722256

**Authors:** Magda Delorence Lugon, Francine Alves Nogueira de Almeida, Pablo Viana Oliveira, Karolinni Bianchi Britto, Pedro Henrique Dias dos Santos, Rafaela Campostrini Forzza, Mário Augusto Gonçalves Jardim, Greiciane Gaburro Paneto

## Abstract

Authentication of açaí products is increasingly important due to the risk of species substitution among morphologically similar *Euterpe* taxa, with implications for food quality, labeling accuracy, and consumer trust. Despite advances in molecular methods, rapid and cost-effective tools for discriminating closely related *Euterpe* species in processed commercial matrices remain limited.

This study evaluated High-Resolution Melting (HRM) analysis targeting two complementary chloroplast markers — psbK-I and ycf1b — as a practical approach for species-level authentication of açaí *(Euterpe oleracea* and *E. precatoria*) and juçara (*E. edulis*) products. In silico specificity analysis confirmed that the ycf1b primer pair shows amplification restricted to the Arecaceae family, supporting the analytical robustness of the method. The combined markers enabled reliable differentiation of all target species, including closely related taxa, with a detection limit of approximately 10% in admixed samples.

When applied to 50 commercial products, HRM successfully authenticated 46 samples, substantially outperforming DNA sequencing, which was limited by amplification failure and mixed chromatograms. Mislabeling was detected in one açaí sorbet and three frozen açaí pulps marketed as açaí but molecularly identified as *E. edulis*, constituting a violation of Brazilian food labeling regulations.

These findings demonstrate that HRM analysis provides a robust, rapid, and scalable strategy for routine species authentication in processed plant-based matrices, with potential for integration into food quality control workflows and large-scale commercial monitoring programs.

## 1. Introduction

Açaí has gained increasing global attention due to its recognized nutritional and functional properties, which have driven the expansion of a wide range of commercial products, including pulps, beverages, sorbets, and dietary supplements (Schauss et al., 2006; de Lima Yamaguchi et l., 2015). Açaí is primarily obtained from *Euterpe oleracea* and, to a lesser extent, from *Euterpe precatoria*, both native to the Amazon region (Oliveira & Schwartz, 2018). In contrast, juçara (*Euterpe edulis*) is a morphologically similar species that has experienced local population declines due to intense exploitation for palm heart extraction, raising conservation concerns at the regional level, particularly within the Brazilian Atlantic Forest (Bourscheid et al., 2011).

Despite their morphological similarity and overlapping physicochemical characteristics, açaí and juçara differ in ecological, nutritional, and conservation aspects, and therefore cannot be regarded as commercially interchangeable (Schulz et al., 2016; Teixeira et al., 2024; Brazil, 2018). The high commercial value of açaí, combined with increasing consumer demand, creates favorable conditions for economically motivated adulteration or substitution, particularly in processed products such as pulps and derived products, where morphological identification is no longer feasible.

Conventional analytical approaches based on physicochemical profiling have been employed to differentiate these species; however, such methods may lack specificity when dealing with closely related taxa and processed matrices (Santos et al., 2014; Schulz et al., 2016). In this context, DNA-based techniques have emerged as reliable tools for food authentication, offering high specificity and sensitivity even in complex and processed samples (Galimberti et al., 2013).

Among these approaches, High-Resolution Melting (HRM) analysis has gained attention as a rapid, cost-effective, and closed-tube technique capable of detecting sequence variations without the need for post-PCR processing (Druml & Cichna-Markl, 2014). HRM has been successfully applied for species discrimination in several food products, demonstrating its potential for routine authenticity testing (Costa et al., 2016). However, its application to differentiate Euterpe species in commercial açaí products remains limited, particularly under real market conditions involving processed samples and potential DNA degradation (Lugon et al., 2021).

Despite the availability of molecular tools for species identification, there is still a lack of rapid, cost-effective, and easily applicable methods capable of discriminating closely related Euterpe pecies in processed commercial products marketed as açaí. This limitation is particularly critical considering that such products are commonly perceived by consumers as originating from the Amazon region, which may not correspond to their actual botanical identity, thereby raising concerns regarding food fraud, mislabeling, and regulatory compliance.

Therefore, this study aims to evaluate the applicability of HRM analysis as a rapid and reliable tool for the authentication of açaí and juçara in commercial products, contributing to the development of molecular strategies to detect species substitution and support food control and regulatory actions.

## 2. Materials and Methods

### 2.1. Samples

The same 50 commercial samples previously analyzed by DNA sequencing in Lugon et al. (2021) were used in this study, enabling a direct methodological comparison between HRM analysis and DNA sequencing under identical sampling conditions. These comprised 47 samples marketed as açaí — purchased from supermarkets, sorbet shops, açaí shops, local fairs, and natural product stores across various regions of Brazil — and three samples sold as juçara, obtained from local fairs, totaling 50 commercial samples. No ethical approval was required for the analysis of commercially available products.

Prior to HRM assay development, nine barcode regions were evaluated in 40 reference samples comprising 19 specimens of *E. edulis*, 15 specimens of *E. oleracea*, and 6 specimens of *E. precatoria* from geographically distinct localities across Brazil, as described in Lugon et al. (2021). Sequence analysis of the psbK-I and ycf1b regions revealed highly conserved intraspecific profiles across all specimens, with interspecific polymorphic sites remaining consistent regardless of geographic origin (Lugon et al., 2021; Fig. S1). Based on this intraspecific sequence conservation, three representative specimens per species were selected for HRM assay development and validation: three specimens of *E. edulis* (vouchers: VIES 24868, VIES 25501, RB 788747), three specimens of *E. oleracea* (vouchers: RB 789434, MG 204258, RB 789437), and three specimens of *E. precatoria* (vouchers: RB 788749, RB 796836, RB 759224). Samples labeled “VIES” were obtained from the herbarium of the Federal University of Espírito Santo, “RB” from the Rio de Janeiro Botanical Garden, and “MG” from the Paraense Emílio Goeldi Museum. Morphological identification was performed based on the descriptions provided by Lorenzi et al. (2010) and/or confirmed by Dr. Rafaela Campostrini Forzza, a taxonomist specializing in Arecaceae at the Rio de Janeiro Botanical Garden.

### 2.2. Marker selection and primer design

The selection of the psbK-I and ycf1b markers was based on previous work by our group, in which multiple barcode regions were evaluated for the authentication of *Euterpe* species in commercial products. In that study, psbK-I showed the best discriminatory performance among the tested loci, supporting its selection for HRM assay development (Lugon et al., 2021). The ycf1b marker was selected as a complementary locus to overcome the limited resolution of psbK-I, as it enabled discrimination between species that exhibited overlapping melting profiles with the primary marker.

Reference sequences were aligned using ClustalW in the BioEdit software (Hall, 1999). HRM primers were designed with Primer3web (Koressaar & Remm, 2007), targeting a 101 bp region containing a single polymorphic variation for psbK-I and a 120 bp region containing two polymorphic variations for ycf1b (Table 1). To predict the melting temperature of each amplicon, version 4.0 of the uMelt DNA Melting Curve Prediction Software (Dwight et al., 2011) was used, enabling species identification based on temperature variation.

**Table 1.**
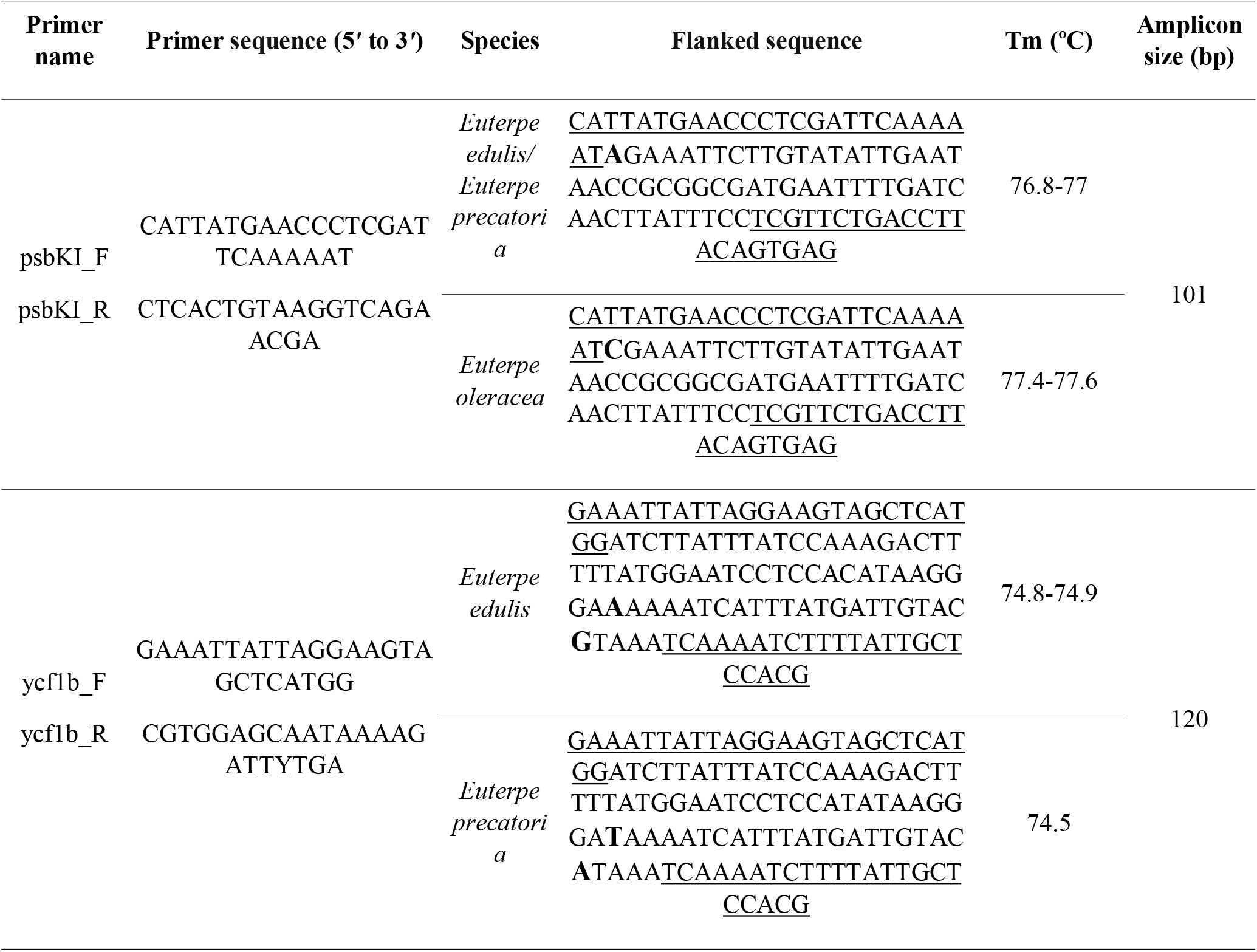
Primers used for HRM analysis targeting the psbK-*I* and ycf1b regions in *Euterpe* species. Flanking sequences are shown, with primer binding sites underlined and SNP positions indicated in bold. Melting temperature (Tm) values were obtained using a LightCycler®96 realtime PCR instrument (Roche Diagnostics).

In silico specificity analysis using Primer-BLAST (Ye et al., 2012) against the NCBI nucleotide database restricted to Viridiplantae (taxid: 33090) confirmed amplification of the psbK-I region across a broad range of Arecaceae species, consistent with the conserved nature of this chloroplast intergenic spacer. Non-palm species commonly present in açaí commercial products, including Musa spp. and *Paullinia cupana*, showed predicted amplification only under reduced primer stringency. In contrast, the ycf1b primer pair showed predicted amplification restricted to the Arecaceae family, with all non-target palm species presenting one or more mismatches in the reverse primer, suggesting substantially reduced amplification efficiency relative to the target Euterpe species. These results support the complementary use of both markers, with ycf1b providing higher taxonomic specificity within the Arecaceae.

### 2.3. DNA extraction, PCR amplification, and sequencing

Leaves from reference samples were macerated in liquid nitrogen to obtain a fine powder, and 30 mg of each sample was incubated with 1 mL of extraction buffer (300 mM TRIS-HCl, pH 8.0; 25 mM EDTA, pH 8.0; 2 M NaCl; 2% (w/v) soluble PVP, MW 40,000; 2% (w/v) CTAB; and 2% (v/v) β-mercaptoethanol). The buffer was preheated to 65 °C, and incubation was carried out for 30 min with periodic manual mixing every 10 min. DNA was then extracted following a modified CTAB protocol (Japelaghi et al., 2011).

The same procedure was applied to 300 μL of commercial samples or 30 mg of açaí powder. Extracted DNA was quantified using a NanoDrop spectrophotometer (Thermo Fisher Scientific Inc., Waltham, MA, USA) and stored at −30 °C for subsequent analyses. PCR amplification and sequencing of all samples were performed as described by Lugon et al. (2021) for comparison purposes.

### 2.4. PCR-HRM analysis

HRM with pre-amplification was performed on a LightCycler® 96 real-time PCR instrument (Roche Diagnostics). PCR reactions were prepared using the EvaGreen® SsoFast− PCR supermix kit, consisting of 5 µL of supermix, 0.5 µL (500 nM) of each forward and reverse primer (Table 1), 10 ng of DNA, and ultrapure DNase- and RNase-free water to a final reaction volume of 10 µL. DNA amplification was conducted in technical triplicate for each sample, including a negative control. For reference specimens, three biologically independent individuals per species were analyzed to assess intraspecific consistency of HRM profiles. Thermal cycling conditions were as follows: initial denaturation at 95 °C for 2 min, followed by 40 cycles of 95 °C for 30 s, annealing at 60 °C for psbK-I primers or 62 °C for ycf1b primers for 30 s, and extension at 72 °C for 30 s.

Following amplification, PCR products were denatured at 95 °C for 1 min and renatured at 40 °C to form DNA duplexes. Melting curves were generated by heating from 65 °C to 97 °C at an increment of 2.2 °C/s, with 25 data acquisitions per degree.

Melting temperature (Tm) data were statistically analyzed using R software version 4.4.3 (R Core Team, 2025) to calculate the mean, standard deviation, and confidence interval for each sample run in triplicate (Tables 2 and 3).

**Table 2.** Melting temperature (Tm) values (mean ± SD and 95% confidence interval) obtained from HRM analysis using the psbK-I marker (101 bp amplicon). NRC = no reference cluster.

| N | Commercial products | HRM cluster | T<sub>m</sub> (°C) | SD | CI (95%) |
| --- | --- | --- | --- | --- | --- |
| 1 | Açai capsule 52 | NRC | - | - | - |
| 2 | Açai sorbet 36 | *E. edulis/E. precatoria* | 76.84 | 0.09 | 76.84±0.10 |
| 3 | Açai sorbet 37 | *E. oleracea* | 77.53 | 0.07 | 77.53±0.08 |
| 4 | Açai sorbet 38 | *E. oleracea* | 77.57 | 0.07 | 77.57±0.08 |
| 5 | Açai sorbet 39 | *E. oleracea* | 77.58 | 0.06 | 77.58±0.07 |
| 6 | Açai sorbet 43 | *E. oleracea* | 77.59 | 0.04 | 77.59±0.05 |
| 7 | Açai sorbet 44 | *E. oleracea* | 77.54 | 0.07 | 77.54±0.08 |
| 8 | Açai sorbet 45 | *E. oleracea* | 77.50 | 0.03 | 77.50±0.03 |
| 9 | Açaí sorbet 58 | *E. oleracea* | 77.48 | 0.02 | 77.48±0.02 |
| 10 | Açaí sorbet 60 | *E. oleracea* | 77.49 | 0.01 | 77.49±0.01 |
| 11 | Açaí sorbet 82 | *E. oleracea* | 77.51 | 0.03 | 77.51±0.04 |
| 12 | Açaí sorbet 86 | *E. oleracea* | 77.53 | 0.02 | 77.53±0.02 |
| 13 | Açaí sorbet 87 | *E. oleracea* | 77.48 | 0.02 | 77.48±0.02 |
| 14 | Açaí sorbet 89 | *E. oleracea* | 77.49 | 0.02 | 77.49±0.02 |
| 15 | Açaí popsicle 56 | *E. oleracea* | 77.57 | 0.03 | 77.57±0.03 |
| 16 | Açaí popsicle 59 | *E. oleracea* | 77.57 | 0.03 | 77.57±0.03 |
| 17 | Açaí popsicle 88 | *E. oleracea* | 77.44 | 0.07 | 77.44±0.08 |
| 18 | Açaí popsicle 91 | *E. oleracea* | 77.59 | 0.05 | 77.59±0.05 |
| 19 | Açaí powder 42 | NRC | - | - | - |
| 20 | Açaí powder 50 | *E. oleracea* | 77.69 | 0.18 | 77.69±0.20 |
| 21 | Açaí powder 51 | *E. edulis/E. precatoria* | 77.43 | 0.11 | 77.43±0.13 |
| 22 | Açaí powder 57 | *E. edulis/E. precatoria* | 76.94 | 0.06 | 76.94±0.07 |
| 23 | Mixed frozen Açaí pulp 47 | NRC | - | - | - |
| 24 | Mixed frozen Açaí pulp 48 | *E. oleracea* | 77.51 | 0.04 | 77.51±0.05 |
| 25 | Mixed frozen Açaí pulp 49 | *E. oleracea* | 77.47 | 0.06 | 77.47±0.07 |
| 26 | Pure frozen Açaí pulp 200 | *E. edulis/E. precatoria* | 76.99 | 0.08 | 76.99±0.08 |
| 27 | Pure frozen Açaí pulp 201 | *E. oleracea* | 77.57 | 0.04 | 77.57±0.04 |
| 28 | Pure frozen Açaí pulp 202 | *E. oleracea* | 77.59 | 0.04 | 77.59±0.05 |
| 29 | Pure frozen Açaí pulp 203 | *E. oleracea* | 77.58 | 0.04 | 77.58±0.05 |
| 30 | Pure frozen Açaí pulp 204 | *E. edulis/E. precatoria* | 76.91 | 0.07 | 76.91±0.08 |
| 31 | Pure frozen Açaí pulp 13 | *E. oleracea* | 77.59 | 0.04 | 77.59±0.04 |
| 32 | Pure frozen Açaí pulp 205 | *E. oleracea* | 77.58 | 0.03 | 77.58±0.03 |
| 33 | Pure frozen Açaí pulp 30 | *E. edulis/E. precatoria* | 76.99 | 0.07 | 76.99±0.08 |
| 34 | Pure frozen Açaí pulp 31 | *E. oleracea* | 77.14 | 0.57 | 77.14±0.65 |
| 35 | Pure frozen Açaí pulp 32 | *E. oleracea* | 77.48 | 0.08 | 77.48±0.09 |
| 36 | Pure frozen Açaí pulp 33 | *E. oleracea* | 77.53 | 0.03 | 77.53±0.04 |
| 37 | Pure frozen Açaí pulp 34 | *E. edulis/E. precatoria* | 76.90 | 0.09 | 76.90±0.10 |
| 38 | Pure frozen Açaí pulp 35 | NRC | - | - | - |
| 39 | Pure frozen Açaí pulp 46 | *E. oleracea* | 77.58 | 0.03 | 77.58±0.03 |
| 40 | Pure frozen Açaí pulp 53 | *E. oleracea* | 77.46 | 0.02 | 77.46±0.02 |
| 41 | Pure frozen Açaí pulp 54 | *E. oleracea* | 77.48 | 0.01 | 77.48±0.01 |
| 42 | Pure frozen Açaí pulp 55 | *E. oleracea* | 77.48 | 0.02 | 77.48±0.01 |
| 43 | Pure frozen Açaí pulp 81 | *E. oleracea* | 77.56 | 0.03 | 77.56±0.03 |
| 44 | Pure frozen Açaí pulp 83 | *E. oleracea* | 77.47 | 0.09 | 77.47±0.09 |
| 45 | Pure frozen Açaí pulp 84 | *E. oleracea* | 77.57 | 0.04 | 77.57±0.05 |
| 46 | Pure frozen Açaí pulp 85 | *E. oleracea* | 77.57 | 0.02 | 77.57±0.02 |
| 47 | Pure frozen Açaí pulp 90 | *E. oleracea* | 77.49 | 0.06 | 77.49±0.06 |
| 48 | Pure frozen Juçara pulp 206 | *E. edulis* | 76.94 | 0.07 | 76.94±0.08 |
| 49 | Pure frozen Juçara pulp 40 | *E. edulis* | 76.89 | 0.16 | 76.89±0.18 |
| 50 | Pure frozen Juçara pulp 41 | *E. edulis* | 76.91 | 0.13 | 76.91±0.16 |

**Table 3.** Melting temperature (Tm) values (mean ± SD and 95% confidence interval) obtained from HRM analysis using the ycf1b marker (120 bp amplicon).

| N | Commercial products | HRM cluster | T<sub>m</sub> (°C) | SD | CI (95%) |
| --- | --- | --- | --- | --- | --- |
| 2 | Açaí sorbet 36 | *E. edulis* | 74.99 | 0.16 | 74.99±0.18 |
| 21 | Açaí powder 51 | *E. precatoria* | 74.53 | 0.05 | 74.53±0.06 |
| 22 | Açaí powder 57 | *E. precatoria* | 74.53 | 0.05 | 74.53±0.06 |
| 26 | Pure frozen Açaí pulp 200 | *E. edulis* | 74.95 | 0.11 | 74.95±0.13 |
| 30 | Pure frozen Açaí pulp 204 | *E. edulis* | 74.85 | 0.09 | 74.85±0.10 |
| 33 | Pure frozen Açaí pulp 30 | *E. precatoria* | 74.50 | 0.05 | 74.50±0.06 |
| 37 | Pure frozen Açaí pulp 34 | *E. edulis* | 74.85 | 0.15 | 74.85±0.17 |
| 48 | Pure frozen Juçara pulp 206 | *E. edulis* | 74.83 | 0.13 | 74.83±0.15 |
| 49 | Pure frozen Juçara pulp 40 | *E. edulis* | 74.84 | 0.13 | 74.84±0.15 |
| 50 | Pure frozen Juçara pulp 41 | *E. edulis* | 74.82 | 0.13 | 74.82±0.15 |

### 2.5. Data analysis

Data analysis was performed using the LightCycler® 96 SW 1.1 software (Roche Diagnostics). Results were evaluated by examining normalized melting curves, difference plots, derivative plots, and melting temperatures (Tm). Commercial samples were classified as authenticated when their HRM profiles clustered within the melting profile range of the corresponding reference species profiles, and as adulterated when clustering with a different species reference group, as determined by the software. Samples that produced no amplification or ambiguous melting profiles were classified as inconclusive.

### 2.6. PCR-HRM validation

The sensitivity of the reaction was assessed using a serial dilution of DNA at concentrations of 10 ng, 1 ng, and 0.1 ng for each reference species with each set of HRM primers. The HRM results were compared to DNA sequencing data to evaluate sensitivity, specificity, precision, and likelihood ratio, following the methodology described by Jilberto et al. (2017).

### 2.7. Mixture detection

To analyze species mixtures using HRM, genomic DNA was extracted from reference samples of *E. edulis* and *E. oleracea*, then diluted to equal concentrations. For admixture preparation, DNA aliquots of *E. edulis* were combined with DNA aliquots of *E. oleracea* in the following ratios: 50:50, 60:40, 70:30, 80:20, and 90:10 for both species. Pure reference samples (100% *E. edulis* and 100% *E. oleracea*) were included as positive controls. HRM results were analyzed using the difference plot, with the 100% *E. edulis* sample serving as the baseline for calculating the melting curves of all other samples.

## 3. Results and discussion

HRM analysis using the psbK-I and ycf1b markers enabled the discrimination of the two species commonly marketed as açaí (*Euterpe oleracea* and *Euterpe precatoria*) and the juçara species (*Euterpe edulis*). The psbK-I amplicon (101 bp) allowed clear differentiation of *E. oleracea* from the other species; however, *E. edulis* and *E. precatoria* showed overlapping melting profiles, indicating limited resolution of this marker for closely related taxa (Fig. 1). In contrast, the ycf1b amplicon (120 bp) successfully resolved this limitation by discriminating *E. edulis* from *E. precatoria*, enabling accurate identification of all target species (Fig. 2). These results highlight the importance of combining complementary markers to achieve reliable species discrimination, particularly in routine authentication workflows where closely related species are involved. As the psbK-I locus is conserved across the Arecaceae family, species discrimination relies on differential HRM melting profiles rather than primer exclusivity, consistent with established approaches in chloroplast-based plant authentication (Ballin et al., 2019; Druml & Cichna-Markl, 2014). The ycf1b primer pair showed higher intrinsic specificity, with non-target Arecaceae species presenting mismatches in the reverse primer, as confirmed by in silico analysis using Primer-BLAST (Ye et al., 2012). The HRM profiles obtained for the three selected reference specimens per species were consistent with the intraspecific sequence conservation previously documented across the broader panel of 40 reference samples from geographically distinct localities in Brazil (Lugon et al., 2021), supporting the representativeness of the selected references for HRM assay development.

**Fig. 1.**
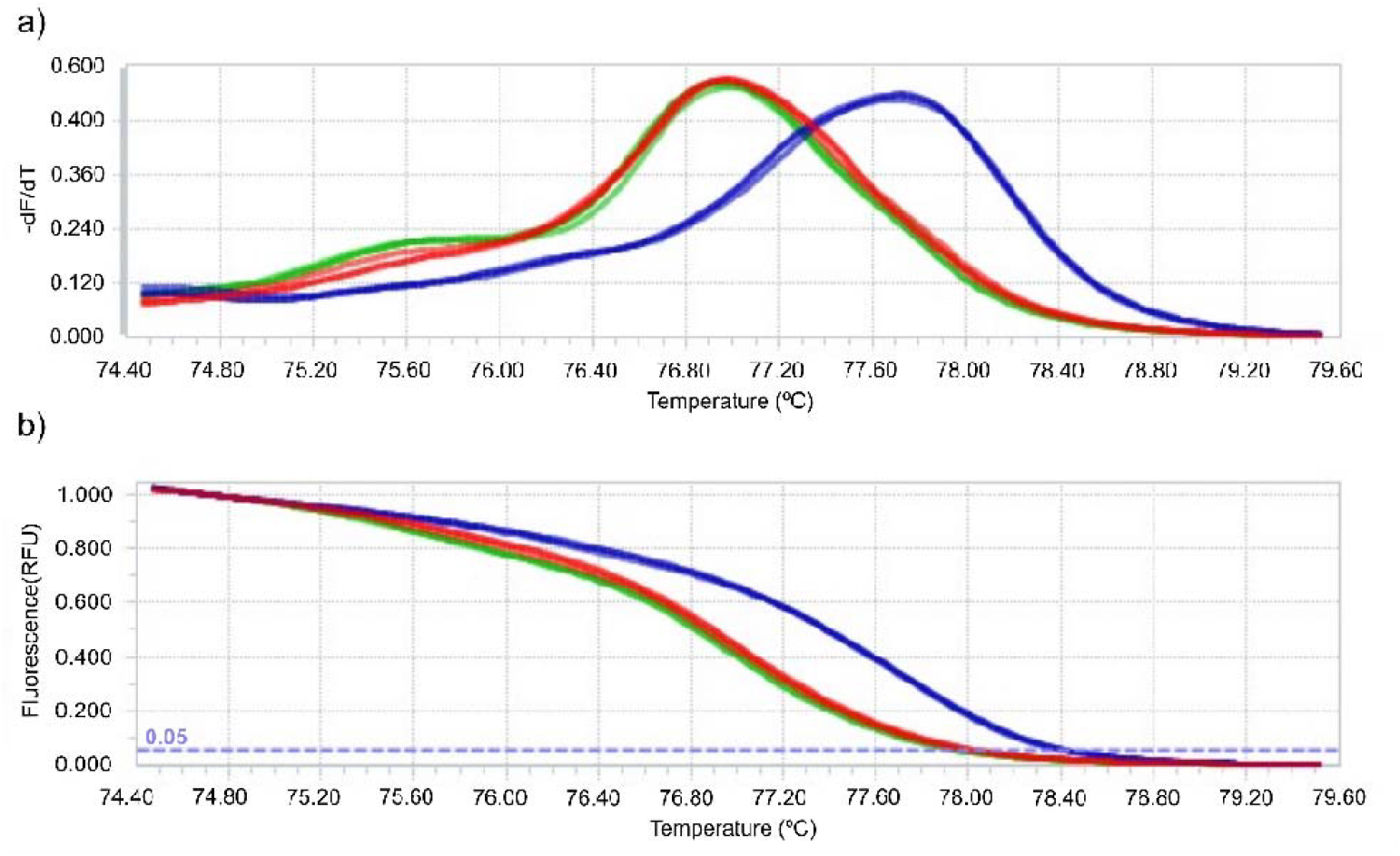
Melting peaks (a) and normalized melting curves (b) of reference samples of *E. edulis* (red), *E. precatoria* (green), and *E. oleracea* (blue) obtained using HRM analysis with psbK-I primers.

**Fig. 2.**
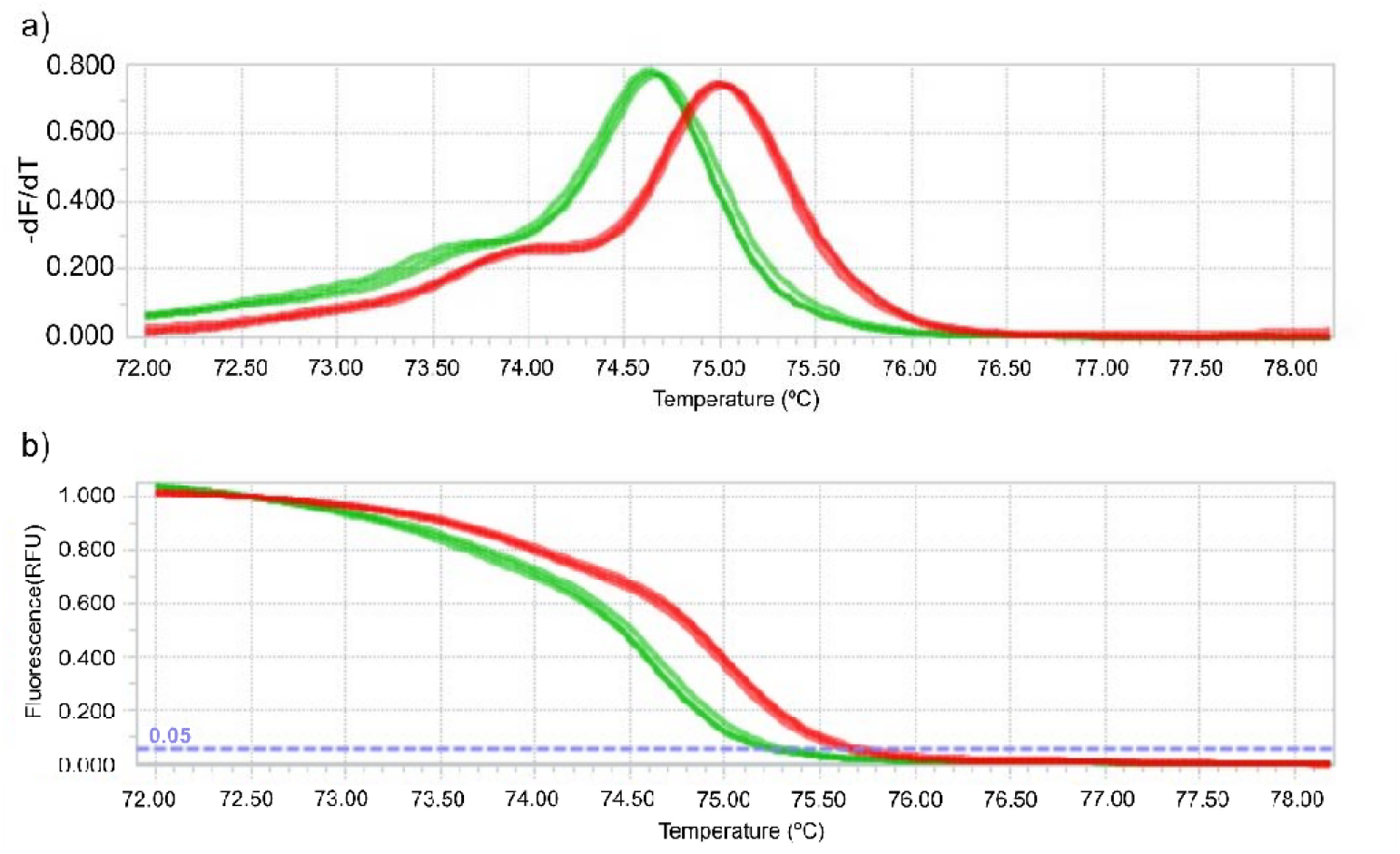
Melting peaks (a) and normalized melting curves (b) of reference samples of *E. precatoria* (green) and *E. edulis* (red) obtained using HRM analysis with ycf1b primers.

The robustness of the HRM assay was assessed using serial DNA dilutions for each reference species with the psbK-I and ycf1b markers. Melting profiles remained consistent across different DNA concentrations, indicating stable amplification behavior and good reproducibility of the assay. In addition, the psbK-I marker generated distinct melting curves across all admixture ratios relative to pure controls (*E. edulis* and *E. oleracea*), enabling the identification of mixed-species samples. The detection limit for both markers was approximately 10%, as 90:10 admixtures were consistently discriminated (Fig. 3), demonstrating the sensitivity of the method for detecting low-level substitution in commercial products.

**Fig. 3.**
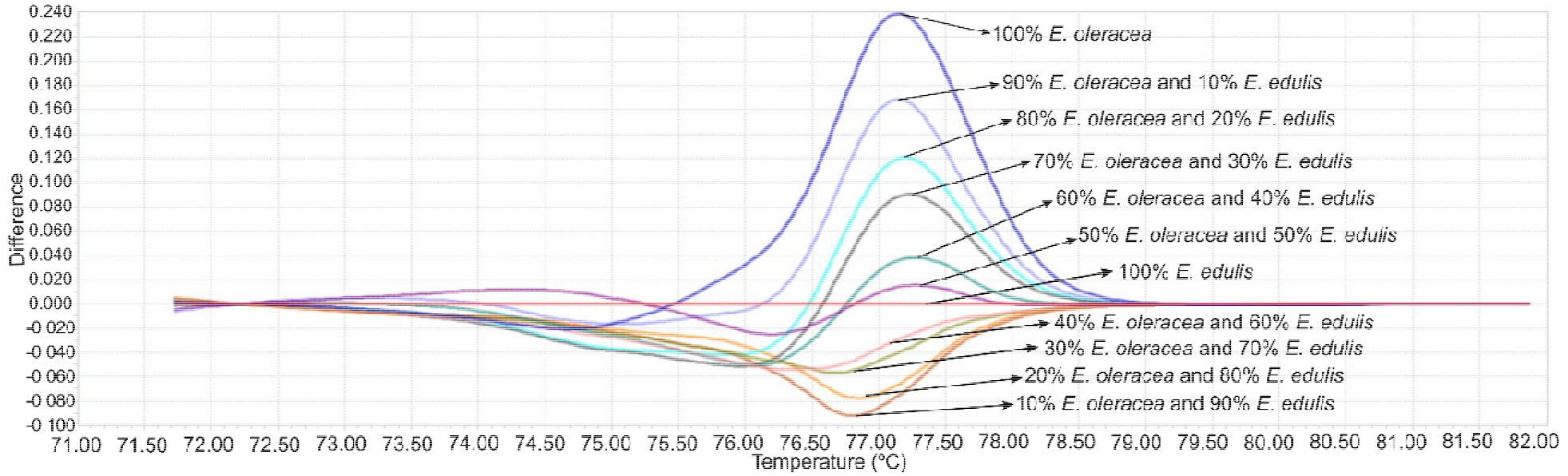
Difference plot obtained using HRM analysis with *psbK-I* primers for reference samples of *E. edulis* and *E. oleracea* at different admixture ratios (50:50, 60:40, 70:30, 80:20, and 90:10). The 100% *E. edulis* sample was used as the baseline for curve normalization.

HRM performance was evaluated against conventional DNA sequencing across 50 commercial samples. While 35 samples produced high-quality sequences, nine showed mixed chromatograms and six failed to amplify, preventing reliable authentication by sequencing. Using the combined psbK-I and ycf1b markers, HRM correctly authenticated 34 of the 35 samples confirmed by sequencing. Importantly, HRM also resolved samples unsuitable for sequencing, successfully authenticating four of the six PCR-negative samples and eight of the nine samples with mixed chromatograms. Overall, HRM enabled the authentication of 46 samples, substantially outperforming sequencing, which authenticated only 35 samples.

Previous studies have reported advantages of HRM over sequencing-based approaches in food authentication. Because HRM targets short DNA fragments, it is particularly effective when analyzing degraded DNA commonly found in processed products. This feature makes HRM well suited for commercial food matrices, where DNA degradation often limits the performance of conventional methods. Consistent with these reports, the present results demonstrate that HRM can overcome key limitations of sequencing, providing a robust and reliable approach for food authentication in challenging commercial samples (Barrias et al., 2025; Grazina et al., 2022).

Using the psbK-I marker, 78.3% of the successfully amplified commercial samples clustered with *E. oleracea* reference profiles and were classified as açaí (Fig. 4, Table 2). However, 21.7% of the samples exhibited overlapping melting curves, preventing discrimination between *E. edulis* and *E. precatoria* using this marker alone. These corresponded to 10 samples, which were subsequently analyzed using the ycf1b marker. Using the ycf1b marker, three samples clustered with *E. precatoria* reference profiles, while seven clustered with *E. edulis* (Fig. 5, Table 3). Among the samples identified as *E. edulis*, three were purchased as juçara and were therefore correctly authenticated. However, the remaining four samples were marketed as açaí but clustered with *E. edulis*, indicating clear cases of mislabeling and potential adulteration. These comprised one açaí sorbet and three pure frozen açaí pulps, consistent with previous findings from the same sample set using DNA sequencing (Lugon et al., 2021). The recurrence of mislabeling in both frozen pulps and sorbet products suggests that species substitution may occur across different product categories and processing conditions, reinforcing the need for authentication methods applicable to a wide range of commercial matrices. Similar concerns regarding species substitution and mislabeling in food products have been widely reported, highlighting the importance of molecular tools for detecting fraud and ensuring product authenticity (Nehal et al., 2021; Haider et al., 2024).

**Fig. 4.**
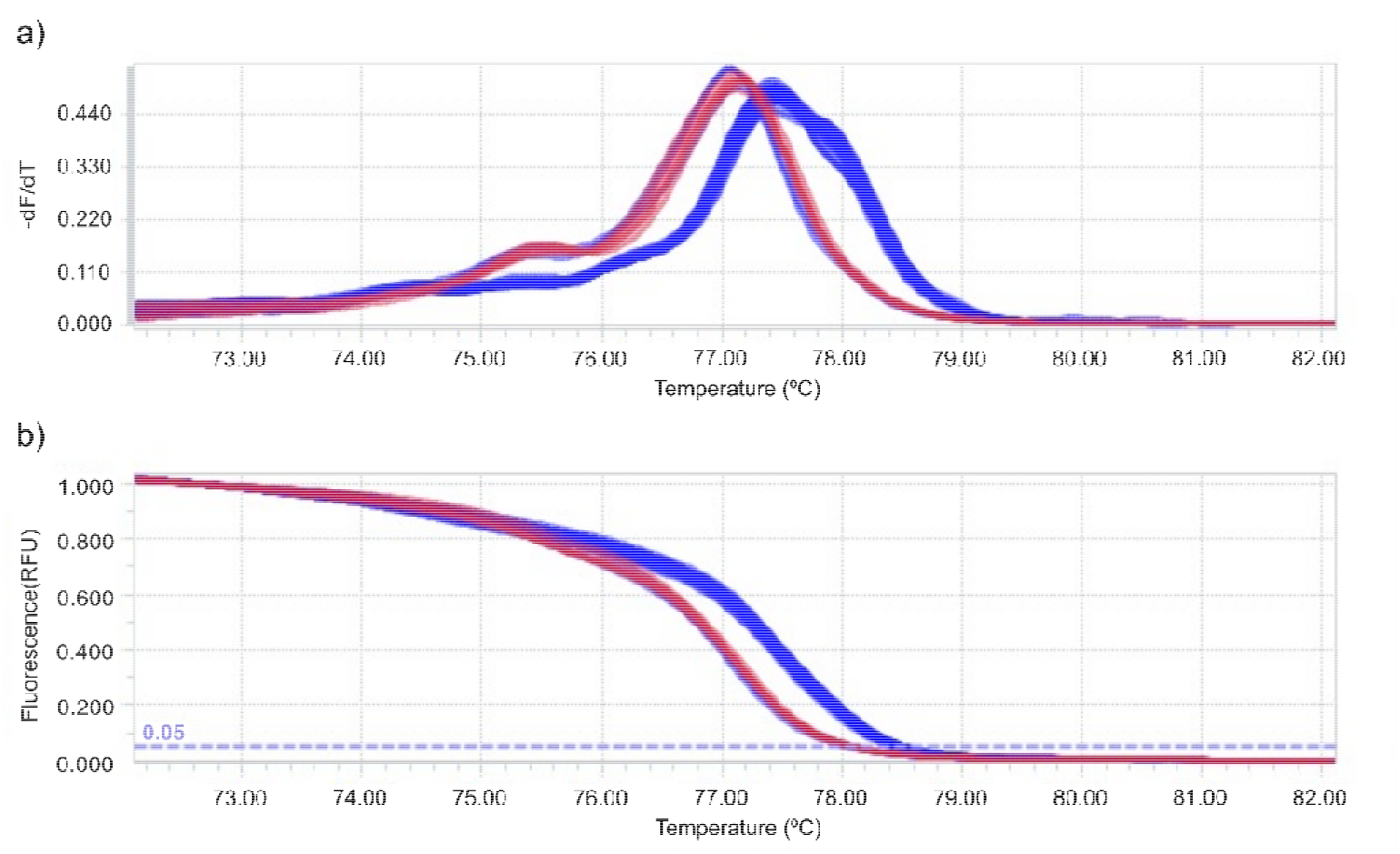
Melting peaks (a) and normalized melting curves (b) of commercial samples compared with reference samples of *E. oleracea* (blue) and *E. edulis* (red) using HRM analysis with psbK-I primers.

**Fig. 5.**
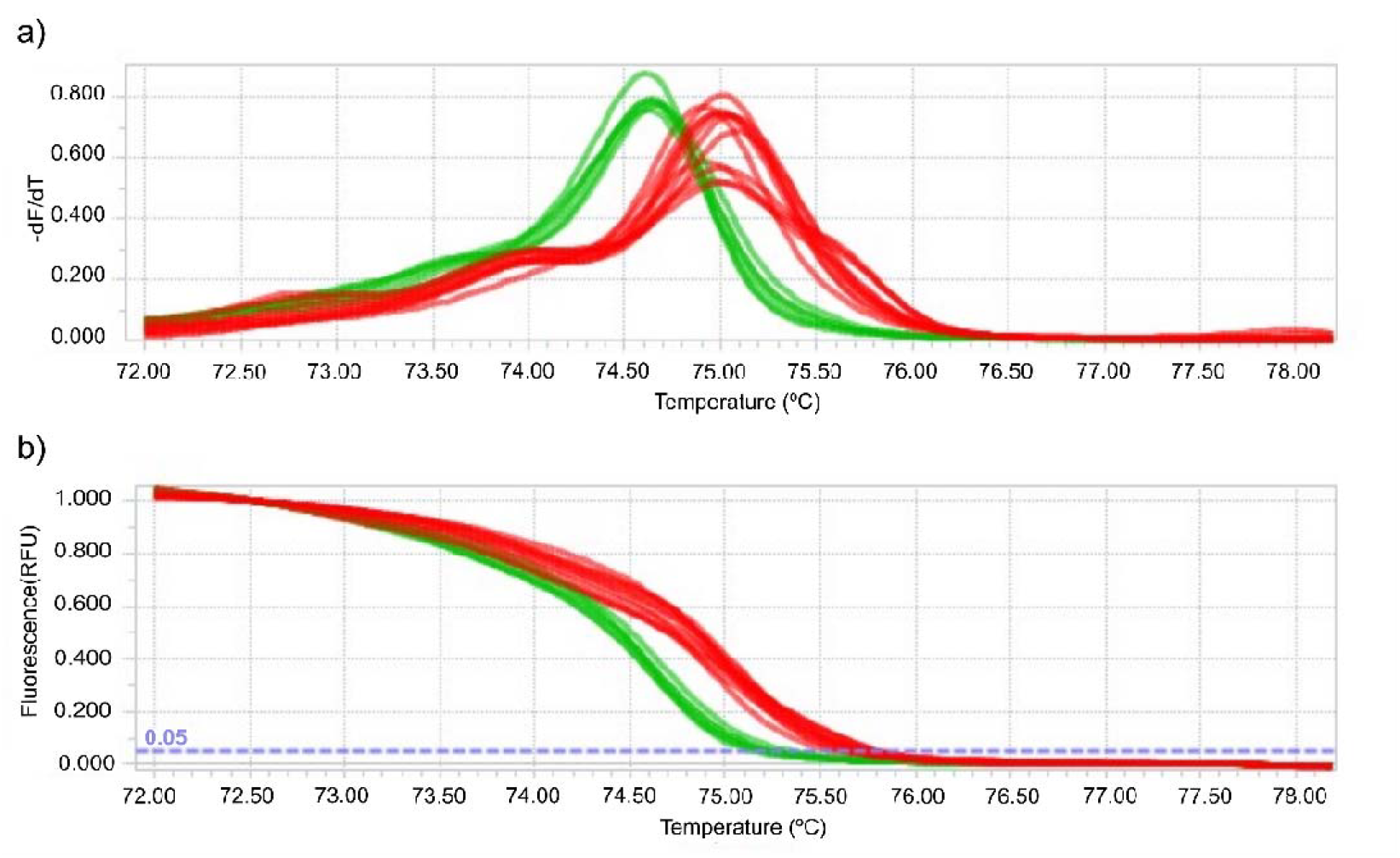
Melting peaks (a) and normalized melting curves (b) of commercial samples compared with reference samples of *E. oleracea* (blue) and *E. edulis* (red) using HRM analysis with ycf1b primers.

The commercialization of juçara pulp has been proposed as a strategy to support the conservation of *Euterpe edulis* and the Atlantic Forest, as fruit harvesting enables seed recovery and contributes to reforestation initiatives. However, the exploitation of this species remains subject to regulatory constraints. These factors, combined with the higher commercial value and market recognition of açaí, may incentivize the mislabeling of juçara-derived products as açaí in informal or illegal markets (Lugon et al., 2021).

From a regulatory standpoint, the mislabeling detected in this study constitutes a direct violation of Brazilian Normative Instruction No. 37/2018 (Brazil, 2018), which establishes that açaí pulp must be derived exclusively from *E. oleracea* or *E. precatoria*, while juçara pulp must originate from *E. edulis*. Furthermore, the standard explicitly prohibits the mixing of pulps from different species without proper declaration on the label. The identification of *E. edulis* in products marketed as açaí therefore represents not only a labeling non-compliance but also a potential case of economically motivated adulteration under Brazilian food legislation. These findings highlight the need for routine molecular monitoring of commercial açaí products by regulatory authorities, and demonstrate that HRM-based methods can provide the analytical basis required for such enforcement actions. The occurrence of samples marketed as açaí but molecularly identified as *E. edulis* in this study supports this concern, highlighting the role of molecular tools in detecting species substitution in commercial products. In this context, HRM provides a practical approach for routine screening, enabling the identification of mislabeling events that may otherwise go unnoticed using conventional methods.

Beyond the direct comparison with sequencing at the sample level, method performance was formally characterized through sensitivity, specificity, and likelihood ratio analyses. The psbK-I marker showed sensitivity values ranging from 96% to 100%, while the ycf1b marker consistently achieved 100% sensitivity, indicating a high capacity for species detection. Both markers exhibited 100% specificity when evaluated against the three target Euterpe species under the experimental conditions tested, demonstrating strong reliability in excluding non-target species within the analyzed sample set. The low negative likelihood ratio (0–4%) further supports the reliability of the method, indicating a high probability that negative results correspond to true absence of the target species. Although the positive likelihood ratio was not defined due to the absence of false positives, the overall results demonstrate high analytical performance. These findings align with recent studies emphasizing the role of DNA-based methods in supporting food control, traceability, and regulatory monitoring (Fontanesi et al., 2024).

The analytical performance of HRM compares favorably with physicochemical approaches previously proposed for the differentiation of *Euterpe* species. Elemental profiling by inductively coupled plasma mass spectrometry combined with linear discriminant analysis has been explored for this purpose; however, its discriminatory power is limited when applied to processed matrices and closely related taxa (Santos et al., 2014). Similarly, anthocyanin characterization and other physicochemical profiling methods may lack the specificity required to distinguish *E. edulis* from Amazonian *Euterpe* species in commercial pulps, particularly when samples have undergone thermal processing or blending (Schulz et al., 2016). In contrast, HRM targets short, species-specific DNA sequences that remain detectable even in degraded samples, offering a more reliable and matrix-independent approach for routine authentication. The combination of high sensitivity, specificity and applicability to processed commercial matrices positions HRM as a practical and analytically superior alternative to physicochemical methods for the authentication of açaí and juçara in the Brazilian market.

Despite these promising findings, some limitations should be considered. The discriminatory power of HRM depends on the selected genetic markers, as demonstrated by the limited resolution of the psbK-I region for discriminating closely related species such as *E. edulis* and *E. precatoria*. The use of a reference panel comprising specimens from geographically distinct localities across Brazil, combined with prior sequence-based validation of intraspecific conservation at the target loci (Lugon et al., 2021), provided a robust basis for HRM assay development and supports the transferability of the method to samples from different regions of the country. In addition, DNA degradation in highly processed products may affect amplification efficiency and melting profile resolution, particularly in samples subjected to intense thermal processing or containing high concentrations of polyphenols and PCR-inhibiting compounds. Another limitation concerns the detection of low-level adulteration below the established threshold (∼10%), which may require further methodological refinement or integration with complementary quantitative techniques, such as digital PCR or next-generation sequencing, to enable more precise estimation of admixture levels. Therefore, HRM should be regarded as a rapid and reliable first-line screening tool, with complementary analytical approaches recommended in cases of ambiguous results or complex sample matrices.

Overall, the present findings reinforce the applicability of HRM as a practical and field-deployable strategy for the authentication of *Euterpe* species in commercial products. Beyond its analytical performance, the method’s speed, low cost, and closed-tube format reduce the risk of cross-contamination, making it particularly suitable for implementation in food control laboratories operating under routine screening conditions. Its broader adoption may contribute to strengthening traceability systems, deterring economically motivated adulteration, and supporting regulatory enforcement in plant-derived food supply chains.

## 4. Conclusion

High-Resolution Melting (HRM) analysis proved to be a rapid, reliable, and cost-effective tool for the authentication of açaí and juçara in commercial pulps and derived products. The combined use of psbK-I and ycf1b markers enabled accurate discrimination among closely related *Euterpe* species, overcoming limitations associated with single-marker approaches. The method demonstrated high sensitivity, specificity, and reproducibility, as well as the ability to detect species substitution in real commercial samples.

Importantly, HRM outperformed conventional DNA sequencing under conditions involving degraded or mixed DNA, highlighting its suitability for processed food matrices. The detection of mislabeling events in products marketed as açaí underscores the relevance of molecular tools in ensuring product authenticity, protecting consumers, and supporting regulatory compliance. Given its speed, cost-effectiveness, and analytical performance, HRM represents a valuable tool for routine screening in food control laboratories. Its application may contribute to strengthening traceability systems and regulatory enforcement. Future studies should focus on improving detection limits for low-level admixtures and integrating HRM with complementary techniques to enable quantitative analysis in complex food matrices.

## Declaration of Generative AI and AI-assisted technologies in the writing process

During the preparation of this work, the authors used ChatGPT (OpenAI) and Claude (Anthropic) to assist with language revision, improvement of textual clarity, and refinement of the scientific writing. After using these tools, the authors critically reviewed and edited all content as necessary and take full responsibility for the integrity, accuracy, and originality of the published article.

## Acknowledgments

The authors express their sincere gratitude to Dr. Márcia Flores da Silva Ferreira for granting permission to use the real-time PCR equipment. This research was funded by the National Council for Scientific and Technological Development (CNPq) (grant number 303420/2016-2) and the Rio de Janeiro Research Foundation (FAPERJ) (grant number E-26/202.778/2018). Additionally, this study was partially financed by the Coordination for the Improvement of Higher Education Personnel (CAPES) under Finance Code 001. The authors are grateful to CAPES for the scholarships awarded to F.A.N. Almeida, K.B. Britto, P.V. Oliveira, and P.H.D. Santos.

## Data availability statement

The DNA sequences generated in the previous study (Lugon et al., 2021) are available in the NCBI GenBank database under the accession numbers listed in Table S1. The HRM profile data generated in this study are available from the corresponding author upon reasonable request.

